# Balancing sensitivity and specificity in distinguishing TCR groups by CDR sequence similarity

**DOI:** 10.1101/526467

**Authors:** Neerja Thakkar, Chris Bailey-Kellogg

**Affiliations:** Department of Computer Science, Dartmouth; Hanover, NH USA

**Keywords:** antigen-specific recognition, CDR classification, immune repertoire, sequence similarity, T cell receptor

## Abstract

Repertoire sequencing is enabling deep explorations into the cellular immune response, including the characterization of commonalities and differences among T cell receptor (TCR) repertoires from different individuals, pathologies, and antigen specificities. In seeking to understand the generality of patterns observed in different groups of TCRs, it is necessary to balance how well each pattern represents the diversity among TCRs from one group (sensitivity) vs. how many TCRs from other groups it also represents (specificity). The variable complementarity determining regions (CDRs), particularly the third CDRs (CDR3s) interact with MHC-presented epitopes from putative antigens, and thus encode the determinants of recognition. We here systematically characterize the predictive power that can be obtained from CDR3 sequences, using representative, readily interpretable methods for evaluating CDR sequence similarity and then clustering and classifying sequences based on similarity. An initial analysis of CDR3s of known structure, clustered by structural similarity, helps calibrate the limits of sequence diversity among CDRs that might have a common mode of interaction with presented epitopes. Subsequent analyses demonstrate that this same range of sequence similarity strikes an appropriate specificity/sensitivity balance in distinguishing twins from non-twins based on overall CDR3 repertoires, classifying CDR3 repertoires by antigen specificity, and distinguishing general pathologies. We conclude that within this fairly broad range of sequence similarity, matching CDR3 sequences are likely to share specificities.

## Introduction

The recognition by T cell receptors (TCRs) of non-self peptide epitopes presented by major histocompatibility complex (MHC) proteins drives the cellular immune response against the non-self offender. In the case of intracellular non-self peptides, e.g., infected or cancerous cells, the ternary peptide-MHC-TCR recognition can lead to the killing of abnormal cells presenting these peptides; in the case of extracellular non-self peptides, e.g., pathogens or biotherapeutics, it can lead to the development of a humoral response to neutralize or clear the antigens containing these peptides. Consequently, modeling and predicting MHC and TCR recognition propensities supports wide-ranging applications, for example: developing vaccines against infectious diseases (1–6) as well as understanding escape mechanisms (7–10), identifying cancer neoantigens and developing specifically targeted vaccines (11–14), discovering potential drivers of allergy, autoimmunity, and tolerance (15–18), and understanding and mitigating anti-biotherapeutic immune responses (19–27).

In the MHC:peptide:TCR recognition process (28), the TCR represents the main source of variability and training in distinguishing of self vs. non-self peptides. MHC is genetically encoded and even the effective diversity across global populations due to allelic variation is limited due to its mode of recognition of the peptide via a fixed binding groove, with degeneracy among the binding groove pockets holding the peptide side-chains (29, 30). In contrast, TCR diversity, produced by VDJ recombination and resulting in hypervariable complementarity determining regions (CDRs), with theoretical diversity estimated to be perhaps 10^15^ (31) and practical diversity in any individual roughly 10^6^ (32–34). An individual’s repertoire diversity is shaped by thymic training against self along with a lifetime of exposure to different antigens, but is presumably much smaller than that of all possible antigens, and there is substantial degeneracy, with different TCRs able to recognize the same antigenic region and the same TCR able to accommodate different antigenic regions (35–38). Thus numerous interesting and important questions center on the relationship between TCR diversity and recognition propensities, including the impacts of genetics vs. training and exposure, the ability of CDRs to accommodate antigenic diversity, and commonalities across pathology- or antigen-specific populations.

The advent of large-scale repertoire sequencing (39, 40), initially for CDRs alone (32, 41–43), and more recently even for paired α/β chains (44, 45), provides opportunities to gain insights into patterns of TCR diversity and recognition. An early study tackled the importance of genetics by studying pairs of monozygous twins, and, among other analyses, found that twin pairs had more identical CDR3 sequences than non-twins (46). More recent publications have shifted the focus from identical sequences to similar sequences. Paired α/β TCRs, resulting from single-cell sequencing and antigen-specific selection, could be classified very well according to their epitope specificities, and the sequences elucidated patterns conferring those specificities (47). Similarly, epitope-specific repertoires from a variety of viral infection contexts pooled across many subjects revealed distinctive motifs, and furthermore the CDRs from a set of *M. tuberculosis* subjects clustered into groups with strong MHC associations enabling design of specific MHC-peptide-TCR interactions (48). An extensive analysis of available TCR sequence data from a wide range of subjects revealed pathogen-specific MHC-TCR associations along with structural insights into MHC and TCR covariation (49).

TCR repertoire sequencing thus provides the opportunity to generalize from a set of samples to patterns than are predictive of relationships among subjects, pathologies, antigens, HLA restrictions, and so forth (47, 49–51). As always in statistical / machine learning approaches, one must thread the needle between under-generalization, missing out on predictions that would be true (i.e., lacking sensitivity), and over-generalization, making predictions that end up not being true (i.e., lacking specificity). Here, we systematically investigate that specificity-sensitivity balance over a number of different datasets and types of predictions, and show that in general there is “sweet spot” in which a range of appropriate trade offs hold, enabling a sufficient number of confident predictions of TCR function from sequence.

## Results

In order to evaluate the relative extent of sequence similarity within and between groups of related CDRs, we study a variety of different CDR sets, using representative approaches based on local sequence similarity, nearest-neighbor classification, and hierarchical clustering. So as to provide a consistent and interpretable basis for drawing conclusions, we focus only on CDR3, using a local alignment score that we term *CDRdist* to assesses similarity (technically dissimilarity, as lower is better), with 0 indicating an exact match and 1 no identifiable similarity. We assess the predictive power of the score by using it to perform nearest-neighbor classification, predicting the group (e.g., associated antigen or disease) to which one CDR belongs based on the known group of the most similar CDR, checking whether the predicted group is correct or not, and thereby evaluating both sensitivity (fraction of CDRs from a group that are correctly predicted to be in that group) and specificity (fraction of CDRs predicted to be in a group that actually are in that group). In order to evaluate only the impacts of similarity, without being confounded by duplicates which can render classification trivial, we do not consider exact matches. In order to gain deeper insights into how the degree of similarity impacts classification performance, we slide a threshold from 0 to 1, making a prediction only if the nearest neighbor is sufficiently similar, and assessing how relaxing the required similarity manifests in specificity vs. sensitivity trade-offs (Fig. 1). The similarity score also provides the basis for clustering, elucidating sequence similarity-based groupings for CDRs and revealing sequence patterns conferring the observed performance trade-offs. We apply these general approaches to characterize several datasets: CDR groups defined by structure, evaluating sequence variation within and between clusters; repertoires from twins, studying the relative similarity between an individual and their twin vs. others; a number of different human and murine repertoires, assessing CDR distinctiveness as it relates to antigen specificity as well as underlying pathology.

**Fig. 1.**
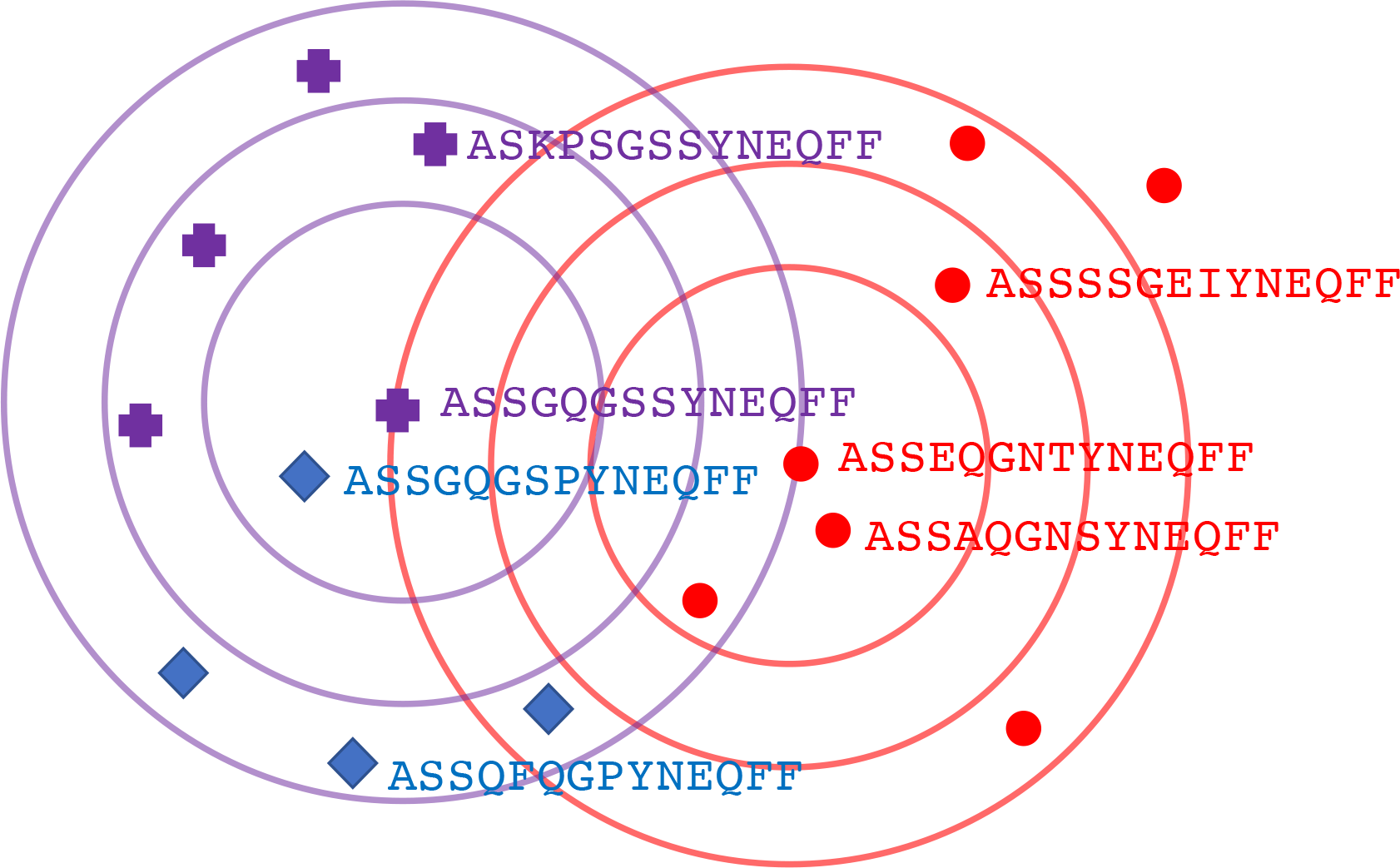
Classifying CDRs by sequence similarity. This illustration plots in a schematic low-dimensional space the locations of CDRs from three classes (red circles, blue diamonds, and purple crosses), and gives sequences for a few. Contour rings show sequence distances of 0.2, 0.3, and 0.4 from an example red CDR and an example purple CDR. The closest CDR to the center red one is also red, so 1-nearest neighbor classification would be correct. In contrast, the closest CDR to the center purple one is actually blue, so 1-nearest neighbor classification would fail. For the red group, increasing the threshold would allow considering more red class members, along with some members from the other classes; one red member lies outside the outermost threshold ring.

### Extent of sequence similarity within/between structural clusters

STCRDab (52) structurally clusters CDR3s into canonical classes, separately for α and β, and separately by length(s). A set of CDR3 sequences and associated structural classes were downloaded and investigated for relative intra- vs. inter-class sequence similarity. After removing duplicates, there were 142 unique sequences across four α groups and six β groups (Tab. 1). While it is common for structural clusters to be characterized by their individual sequence profiles, we also sought to understand the extent to which the sequence pattern from one cluster could be generalized before encroaching on another cluster.

**Table 1.** Unique CDRs from different structural classes according to STCRDab (52).

| **Class name** | **Length(s)** | **Number of unique sequences** |
| --- | --- | --- |
| A3-10-A | 10 | 14 |
| A3-13-B | 13 | 3 |
| A3-13-A | 13 | 6 |
| A3-10_11_12-A | 10 | 10 |
| B3-10_11_12_13-A | 10, 11, 12, 13 | 56 |
| B3-10_11-A | 10, 11 | 20 |
| B3-12-A | 12 | 6 |
| B3-12-B | 12 | 10 |
| B3-13_14-A | 13, 14 | 14 |
| B3-14-A | 14 | 3 |

The distance from each unique CDR to the most similar (but distinct) CDR within its structural class tends to be smaller than the distance to the most similar (but distinct) CDR from another class (Fig. 2(a,b)). When the distance is less than about 0.2 or 0.3, the closest CDR tends to be within the same structural class, while when it is above about 0.6 or 0.7, the CDRs tends to be within different classes. This observation supports the use of nearest-neighbor classification, predicting structural class from sequence matches, which we elaborate to study specificity-sensitivity trade-offs by subjecting it to a similarity threshold; i.e., only make a prediction if the nearest neighbor is closer than a given threshold. When the threshold is less than about 0.2, many CDRs are unclassified but those that are tend to be correct; between 0.2 and 0.4, the number of unclassified CDRs drops substantially while still retaining high specificity; and above that range the specificity drops in order to obtain further sensitivity (Fig. 2(c,d)).

**Fig. 2.**
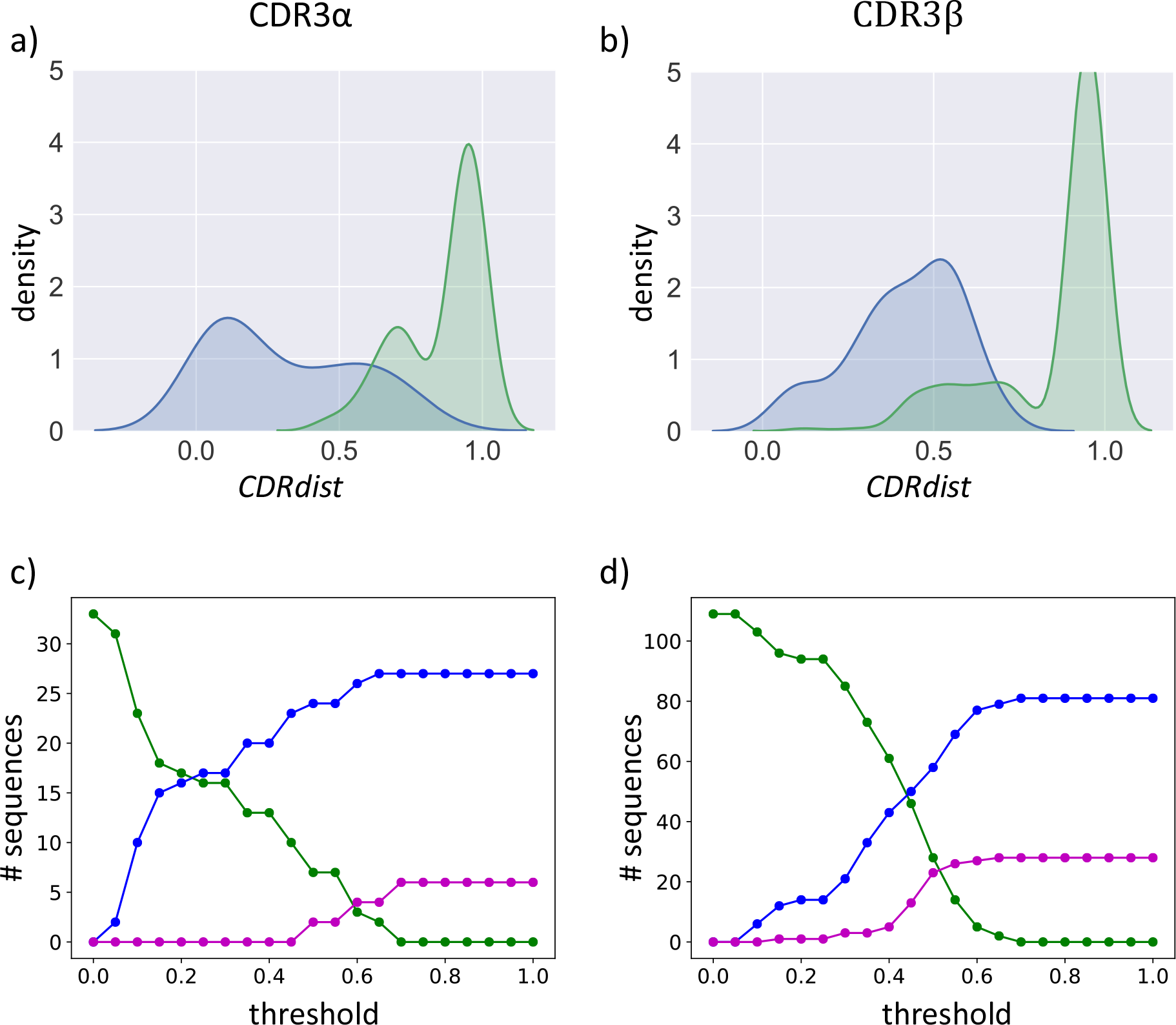
Sequence similarity in STCRDab structural clusters. **(a, b)** Minimum sequence distances within (green) vs. between (blue) structural classes for **(a)** CDR3α sequences and **(b)** CDR3β sequences, plotted as a density estimate. **(c, d)** Accuracy of nearest-neighbor classification using sequence similarity to predict structural cluster for **(c)** CDR3α sequences and **(d)** CDR3β sequences. As the threshold required to make a classification is varied from 0 to 1 (*x*-axis, with 0 indicating identity), the number of sequences (*y*-axis) that are unclassified (green line) decreases, trading off how many are correctly (blue line) vs. incorrectly (magenta) classified.

In order to more directly explore the relationship between sequence similarity and structural similarity, structures were downloaded from STCRDab and the CDR3β loops extracted for those in which electron density was present. When the same sequence was present in multiple structures, a representative was chosen as that with minimal sum of main-chain root mean squared deviation (RMSD) to the others. For each such CDR, the most similar sequence in its STCRDab structural class and the most similar sequence from another class were compared, in terms of both *CDRdist* and RMSD. This comparison (Fig. 3) thus elaborates the implications of Fig. 2, characterizing when a closer (and close enough) sequence implies a closer structure. Some examples are illustrated, limited to cases with good sequence similarity, such that a classification decision would be made under the thresholding approach above. In (a), sequence-based classification is correct, but actually does not yield the best structural match. In (b-d), classification is also correct, and in these cases yields similar (b), better (c), and much better (d) structural matches. In (e) and (f), classification according to STCRDab classes is incorrect but actually consistent with the relative structural similarity: the other structure is more similar (e) or about the same (f); we note however that in (e) neither structure is particularly similar. Finally, in (g) classification is incorrect, and the closest sequence in the same structural class is more similar to that in the other class.

**Fig 3.**
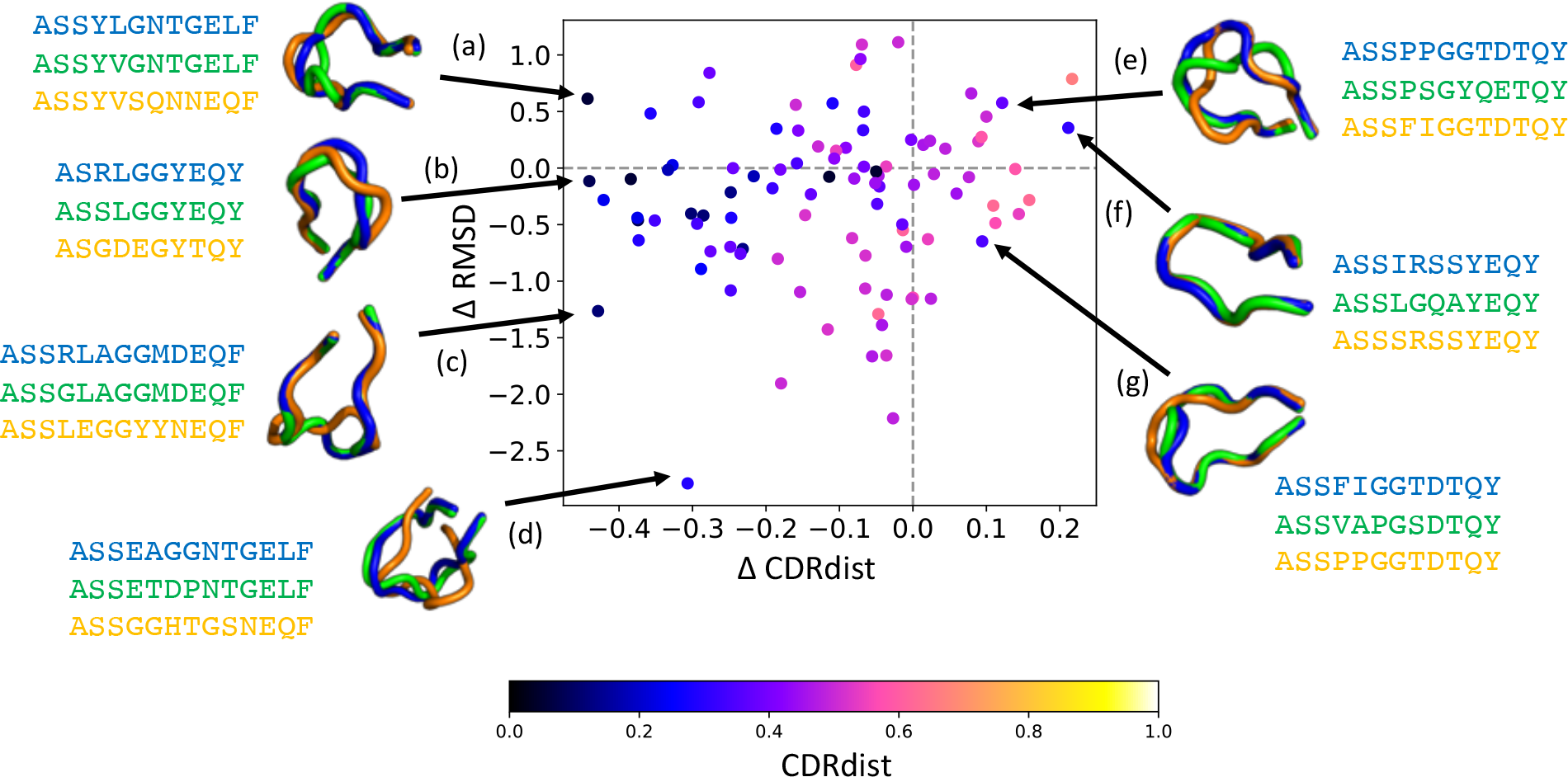
Sequence similarity differences vs. structural similarity differences. Each point represents a CDR3β, and its coordinate represents the difference in sequence score (*x*-axis) and structural similarity (*y*-axis) between the most similar sequence in its STCRDab structural class and the most similar sequence in another STCRDab structural class. A negative number indicates that the CDR is more similar to the neighbor in its structural class than to the other neighbor; i.e., left indicates more similarity in sequence and bottom more similarity in structure to an in-class CDR than to an out-of-class one. Color represents the sequence score between the CDR and the in-class neighbor. Structural superpositions highlight example relationships between a target CDR (blue), its in-class neighbor (green), and its other neighbor (orange).

**Fig. 4.**
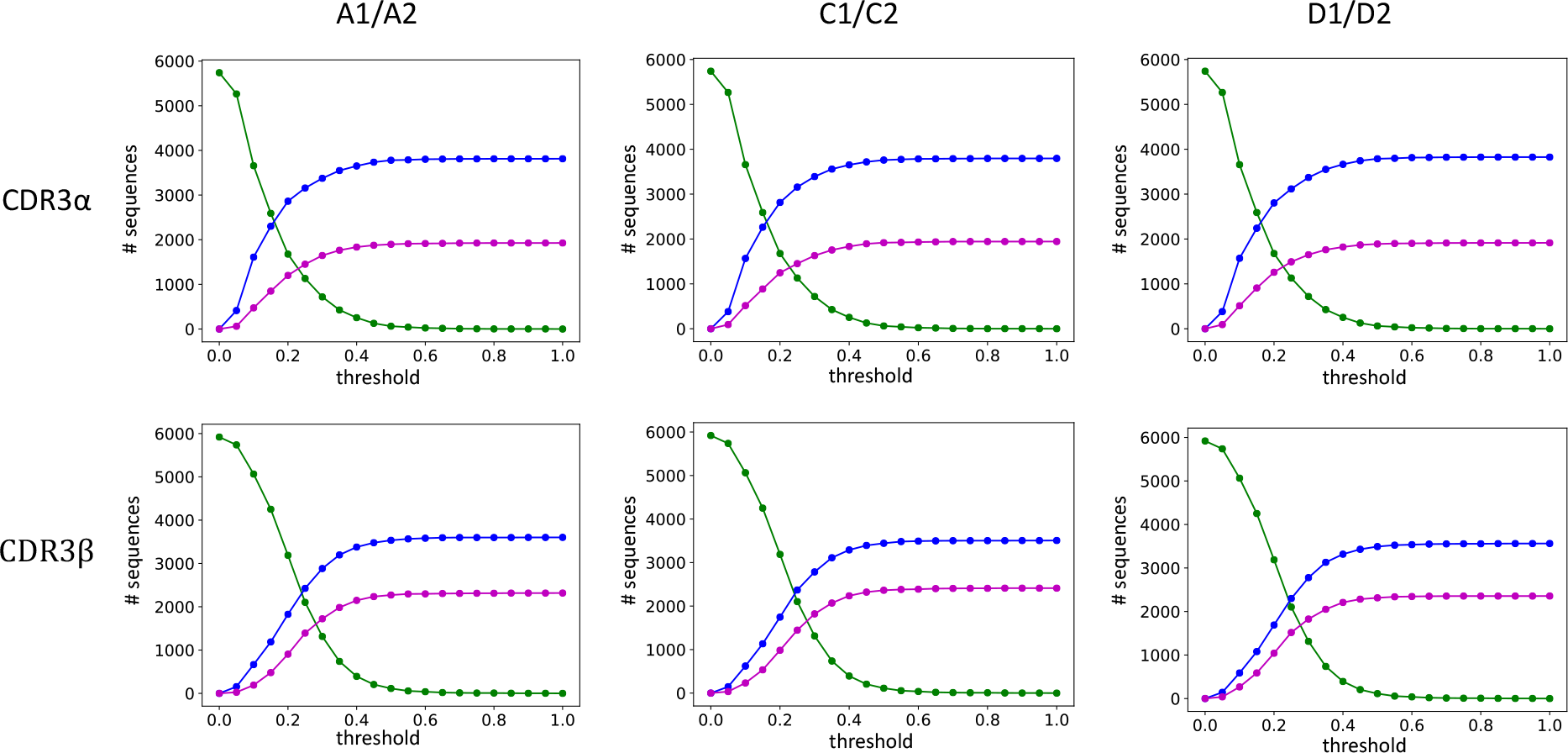
CDR3 similarity in twins vs. non-twins. The plots track the performance of nearest-neighbor classification, using sequence similarity to predict whether a CDR is from a twin or an unrelated individual. As the threshold required to make a classification is varied from 0 to 1 (*x*-axis, with 0 indicating identity), the fraction of sequences (*y*-axis) that are unclassified (green line) decreases, trading off the fraction that are correctly (blue line) vs. incorrectly (magenta) classified.

This analysis helps calibrate the general level of confidence one can have that two CDRs of a given degree of local sequence similarity are likely to adopt similar structures, under a readily interpretable classification approach.

### CDR similarity across repertoires from twins

An early landmark study in TCR repertoire analysis characterized three pairs of monozygous twins, evaluating general characteristics of the repertoires (e.g., diversity) as well as the extent of identity across subjects (46). The investigation revealed that the number of identical CDR3 sequences between two individuals was significantly increased if they were twins. We sought to relax the identification of identical sequences across individuals to allow for different degrees of similarity, in particular to test whether twins had more similar sequences than non-twins. As throughout this paper, we explicitly did not consider exact matches, in this case thereby evaluating the “residual” information beyond the previously studied identity. In order to focus the analysis on the strongest signal, we considered only the 1000 most abundant CDR3 sequences from each repertoire, with read counts ranging from over 100,000 down to around 100. Zvyagin et al. (46) observed that shared clonotypes were significantly higher among the most abundant CDR3β sequences for any pair of individuals, and this was even more evident for twins, so we focused on these more significant sequences.

For each unique CDR from each subject, the closest non-identical CDR in each other subject was identified. These matches served as the basis for evaluating nearest-neighbor classification, assessing whether the closest CDR was from a twin (correct) or a non-twin (incorrect). Over the range of required similarity thresholds, the number of correct classifications outpaces that of incorrect ones (Fig. 3), though not as strongly as for the structural clusters. For CDR3α, at a threshold of 0.2 about 49% of each twin pair’s CDRs are correctly classified and about 21% incorrectly classified, with the remaining 29% unidentified. Raising the threshold to 0.3 yields roughly 59% correct, but at the cost of roughly 29% incorrect, leaving only 12% unidentified.

Further increases in the threshold continue this trend, with 0.4 resulting in 64% correct but 32% incorrect, and 4% unidentified. Results for CDR3β follow the same sensitivity-specificity trend over this range, but at a lower accuracy: at a threshold of 0.2, there are about 30% correct vs. 16% incorrect; at 0.3 the trade-off is 48% correct vs. 30% incorrect, and at 0.4, 56% correct vs. 37% incorrect. Overall, since identical sequences were explicitly excluded from the classification, this result demonstrates that twin repertoires share not only more identical sequences, but also more similar sequences.

To quantify the significance of the difference in twin vs. non-twin nearest neighbors, the distribution of the closest twin distance was compared to that of the closest non-twin match with a Mann-Whitney U test. CDR3β matches between twins in pair A are closer to each other than to their closest non-twin matches (p-value 7 × 10^−4^) as are those in twin pair D (1 × 10^−9^); however, the distinction does not hold for twin pair C (0.24). For CDR3α, however, only twins A are significantly different from others (0.017), with C marginally above a 5% cutoff (0.07) and D insignificant (0.198). Interestingly, the CDR3α classification performance is better than that for CDR3β, even though the overall distributions are more similar, suggesting that focusing specifically on close-enough pairs reveals additional valuable information distinctive of twin pairs.

To explore patterns of similarity and differences within and between twin pairs, CDRs in each individual’s repertoire were clustered according to *CDRdist* at a maximum distance of 0.3. Clusters from the different individuals were then compared according to an aggregate similarity score, *clusterdist*, computed as the sum of cluster member distances. Fig. 5 illustrates some of the patterns of cluster specificities, with motifs revealing the determinants of specificity (or not). In these examples, a C-terminal `TGELFF` appears to be common to some CDRs in all of the individuals, while `DRE***QPQHF` is more specific to individual A1, and `G**YEQYF` distinguishes the twin pair A1/A2 from the other individuals.

**Fig. 5.**
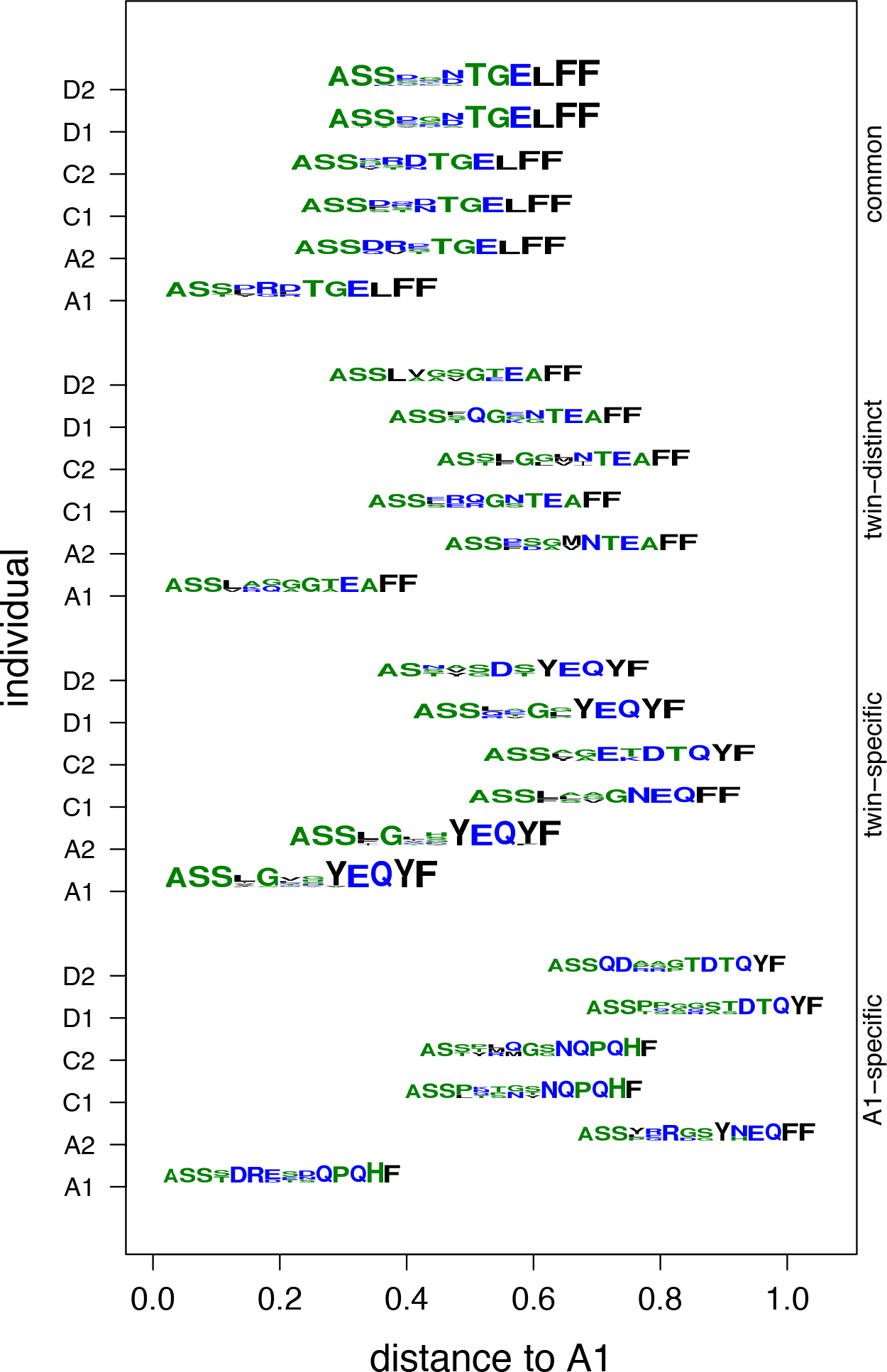
Example twin motifs with different specificity patterns. Motifs represent sequences for four different clusters from subject A1, and the closest clusters to each of those in each of the five other individuals. The *x*-axis indicates the distance from the A1 cluster to a cluster from another individual, assessed as the average over pairs of CDRs. Sets of clusters illustrate different types of patterns: **(common)** all of the clusters are similar to the cluster from A1 (within 0.3); **(twin-distinct)** clusters from unrelated individuals are closer than that from the twin; **(twin-specific)** the twin’s cluster is closer than those from other individuals; **(A1-specific)** the cluster is far from all clusters from all other individuals.

### Classification of epitope specificity by CDR similarity

As discussed in the introduction, a pair of seminal studies published in 2017 studied epitope-specific TCR repertoires across many individuals. This section characterizes sensitivity-specificity trade-offs in predicting within these datasets which epitope each CDR3 recognizes based on the epitopes for similar CDR3s.

Dash et al. (47) used pMHC-tetramer selection and single-cell amplification to collect 4635 paired α/β TCR sequences from 10 epitope-specific repertoires. The 1211 unique CDR3α and 1244 unique CDR3β mouse sequences came from 78 mice and were associated with the epitopes NP, PA, F2, and PB1 (presented during influenza infection) and M38, m139, and M45 (presented during murine cytomegalovirus infection). The 276 unique CDR3β and 294 unique CDR3α human sequences came from 32 humans and were associated with the epitopes M1 from influenza virus, pp65 from human cytomegalovirus, and BMLF1 from Epstein-Barr virus. Among other analyses, a nearest-neighbor classification approach was employed to predict epitope specificity based on a specialized sequence similarity score comparing entire TCR sequences with both α/β chains, using BLOSUM substitution scores with higher weight given to both CDR3 regions. With nearest neighbor classification, 78% (mouse) and 81% (human) of the TCRs were assigned to their correct epitope group. We again sought to characterize how specificity and sensitivity vary, and as throughout the paper directly focused on unpaired CDR3 only and the single most similar sequence to a given one.

Fig. 6 illustrates performance over the range of allowed sequence similarity thresholds. At 0.2, 55% of the mouse CDR3βs are classified correctly, as are 43% of the human ones, with 18% mouse and 5% human incorrect. Relative to the CDRs that are actually identified (73% mouse and 48% human), these fractions are 75% mouse correct and 89% human correct, comparable to the previous results (78% and 81%, respectively). Thus even using just the CDR3β alone, *if the single closest neighbor is close enough* then it is highly predictive of the epitope group. Relaxing the threshold to 0.3 yields more correct classifications, 59% mouse and 53% human, at the cost of more incorrect, 31% mouse and 8% human (8%). Consequently the accuracy among those identified is somewhat lower but still comparable, 66% mouse and 86% human. Further relaxing the threshold continues to yield further increases for both correct and incorrect classifications, e.g., at 0.4, 62% of the mouse sequences are correctly classified vs. 13% incorrectly, and 59% correct vs. 17% incorrect for human, which translates to 63% mouse and 78% human correct among those identified. Thus the suitable balance between accuracy and number of predictions again appears to fall in the 0.2 to 0.4 range as we observed first in the structural clusters as evidence of similar conformation. Similar trends hold for analysis of the CDR3α sequences, but CDR3β sequences had greater predictive power, particularly for murine sequences. Other studies, e.g., (46, 48), have likewise found CDR3β sequences to be more informative than CDR3α sequences.

**Fig 6.**
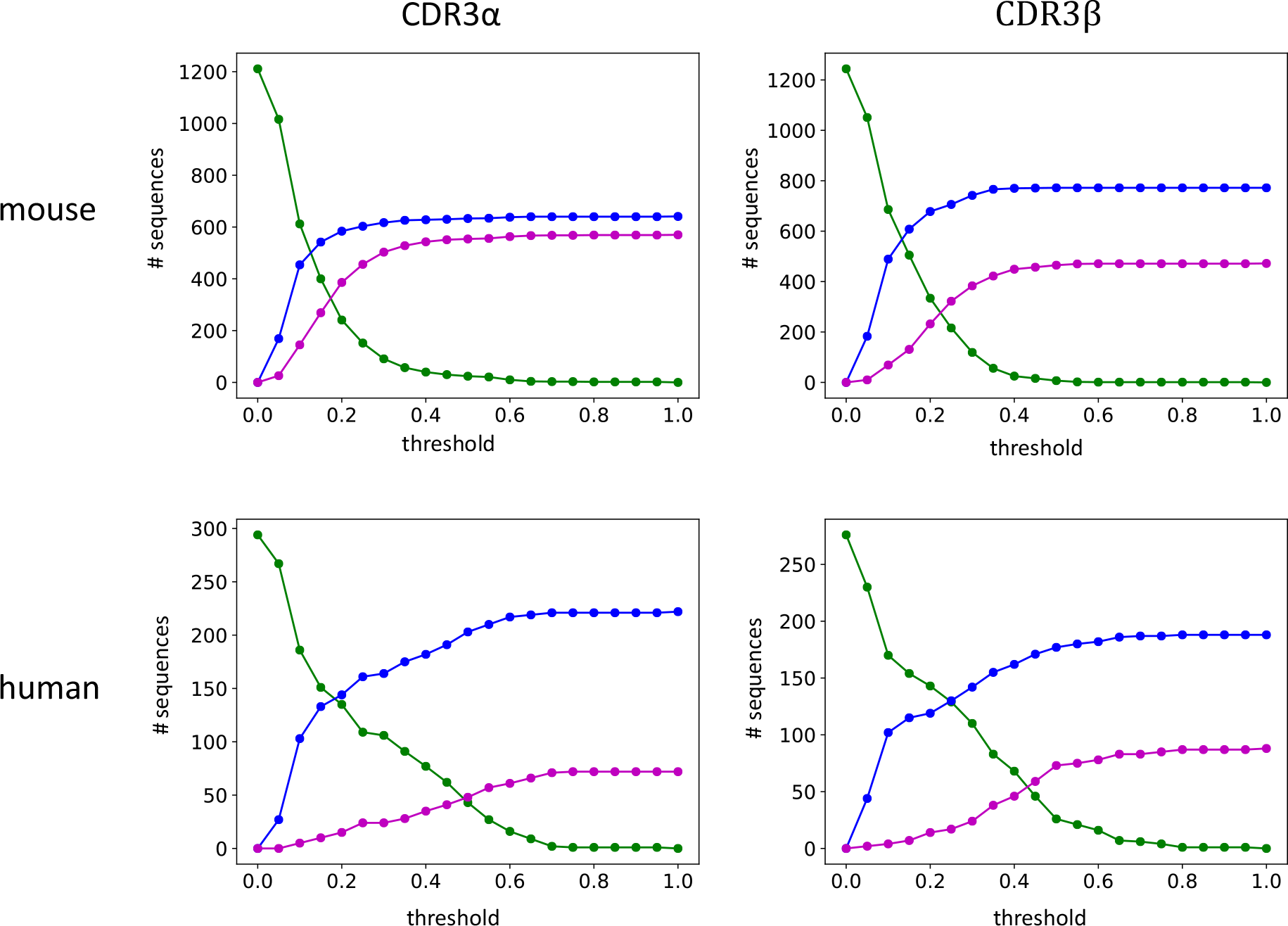
Epitope specificity in Dash et al. repertoires. The plots track the performance of nearest-neighbor classification, using sequence similarity to predict a CDR’s epitope from a set of seven different murine epitopes or from a set of three different human ones. As the threshold required to make a classification is varied from 0 to 1 (*x*-axis, with 0 indicating identity), the fraction of sequences (*y*-axis) that are unclassified (green line) decreases, trading off the fraction that are correctly (blue line) vs. incorrectly (magenta) classified.

Each of the epitope-specific repertoires was clustered, and as with the twins dataset, the clusters were evaluated for relative specificity. Within the murine and human groups of repertoires, the cluster similarity score *clusterdist* was computed for each pair of clusters. Each cluster’s specificity to its repertoire was characterized by the smallest *clusterdist* to a cluster from a different repertoire, since a relatively small *clusterdist* indicates that the sequence pattern defining a cluster common to epitopes in other repertoires while a relatively large score indicates that no cluster in another repertoire has similar epitopes. Fig. 7 illustrates some examples of relatively specific and relatively non-specific clusters at a threshold of 0.3. For example, TCS*GTGG*NYAEQFF is common to both PB1 and M38 from mice, with a distance of only 0.1 between the two clusters. On the other hand, `SCG**GTNEKLFF` is distinct to the human BMLF1 repertoire. Its closest motif is ATGRGG*IGEQYF in p65 at a distance of 0.72, which is relatively large, indicating that `SCG**GTNEKLFF` is unique to BMLF1.

**Fig 7.**
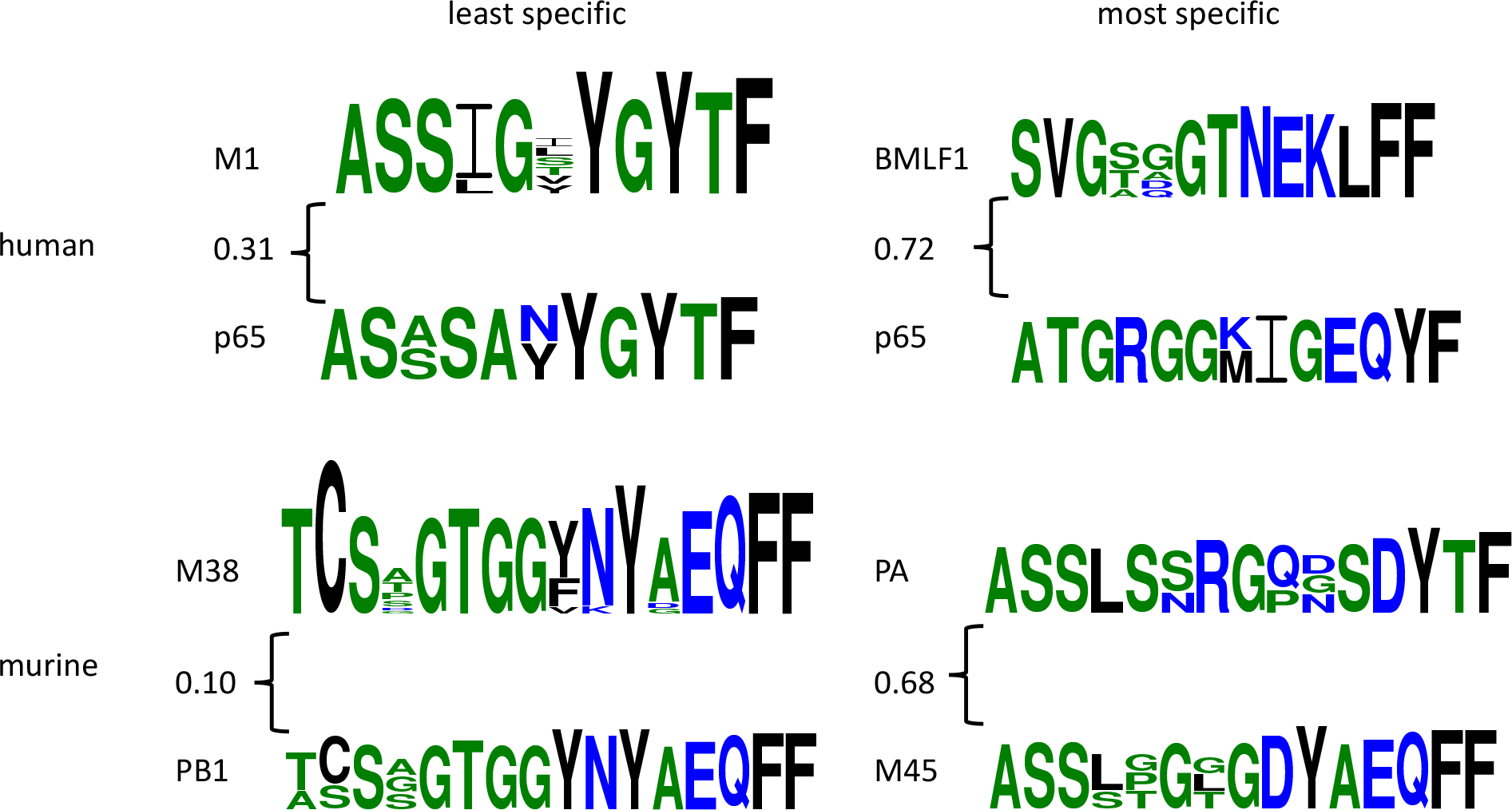
Example relatively specific and relatively non-specific epitope clusters in Dash et al. repertoires. Each motif represents sequences from a cluster of epitopes at a 0.3 threshold. The distance between an example cluster from one cluster and the nearest from another, computed as the average over pairs of their CDRs, is annotated at the curly brace.

As the threshold varies, the resulting clusters evolve, as illustrated in Fig. 8 for some examples. As the threshold increases, clusters tend to become larger, containing more unique sequences, and are therefore more diverse and less specific. In particular, for the NP cluster, the nearest cluster in a different repertoire at 0.2 is at a *clusterdist* of 0.52, but that falls to 0.19 for the 0.3 NP cluster; similarly, for PB1-F2 the nearest other-repertoire cluster at 0.2 is 0.61 away, down to 0.38 at 0.4.

**Fig 8.**
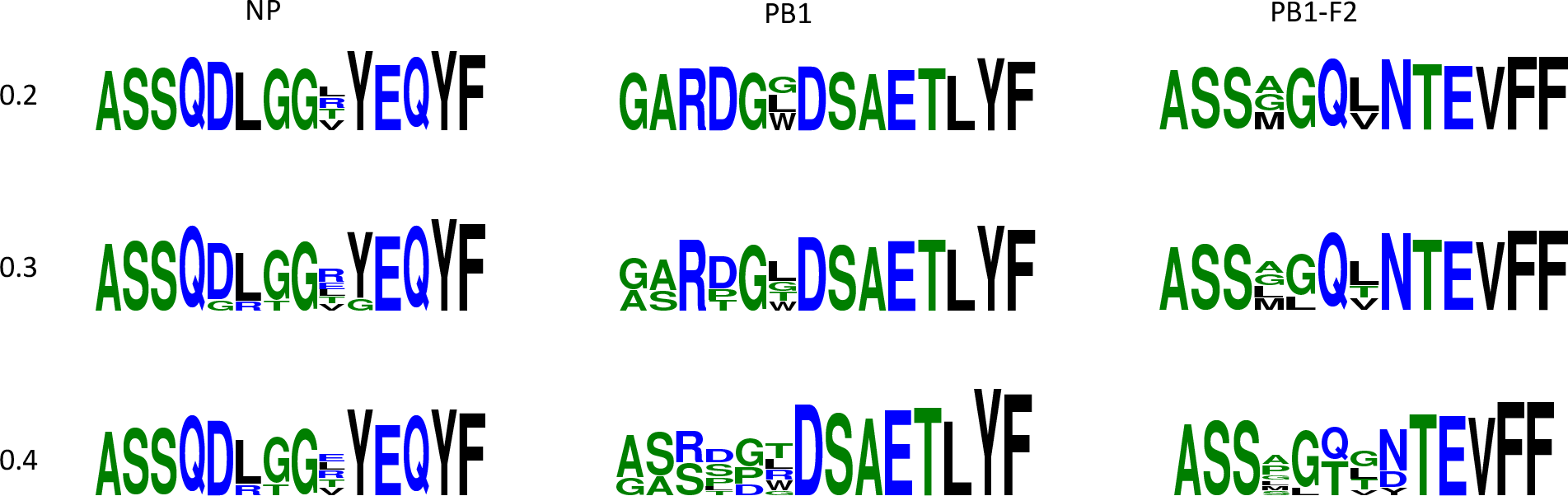
Example progression of clusters in Dash et al. repertoires as the threshold is relaxed. Motifs represent clusters from three different repertoires (columns: NP, PB1, PB1-F2) at three different thresholds (rows: 0.2, 0.3, 0.4).

Turning to the other recent large TCR repertoire study, Glanville et al. (48) collected 2,068 unique sequences using the pMHC tetramers to isolate antigen-specific T cells spanning eight tetramer antigen-HLA specificities: pp50 associated with HLA+A1 (271 sequences), NP177 associated with HLA-B7 (213), pp65 + HLA-A2 (155), pp65 + HLA-B7 (56), BMLF1 + HLA-A2 (700), M1 + HLA-A2 (56 sequences), and NP44 + HLA-A1 (24 sequences). Their analysis of this data showed that the antigen-specific repertoires tended to share more similar sequences, and revealed some 2-, 3-, and 4-mer motifs enriched in different repertoires. We sought to build on this analysis by once again systematically evaluating the extent of generalization and predictiveness supported by sequence patterns.

Nearest-neighbor classification was performed for each pair of specificities at thresholds of 0.2, 0.3, and 0.4, predicting the epitope+HLA of one CDR based on that of the most similar one (Fig. 9). NP44 and pp65 associated with HLA-B7 generally do very well, and BMLF1 also does relatively well. NP177 and pp65 associated with HLA-A2 are more easily confused with other epitopes. On average, over all the pairwise comparisons, a threshold of 0.2 yields about 21% correctly classified vs. 3% incorrectly classified, with 71% unidentified. Raising the threshold to 0.3 yields 42% correct vs. 8% incorrect with 52% unidentified sequences, and 0.4 continues the trend to 59% vs. 16% with 25% unidentified. The standard deviations on these correct percentages are around 11-13%, as some pairs are clearly quite better than others. Overall, these tests confirmed that the 0.2 to 0.4 range balances accuracy and a sufficient number of predictions for the epitopes in this dataset.

**Fig 9.**
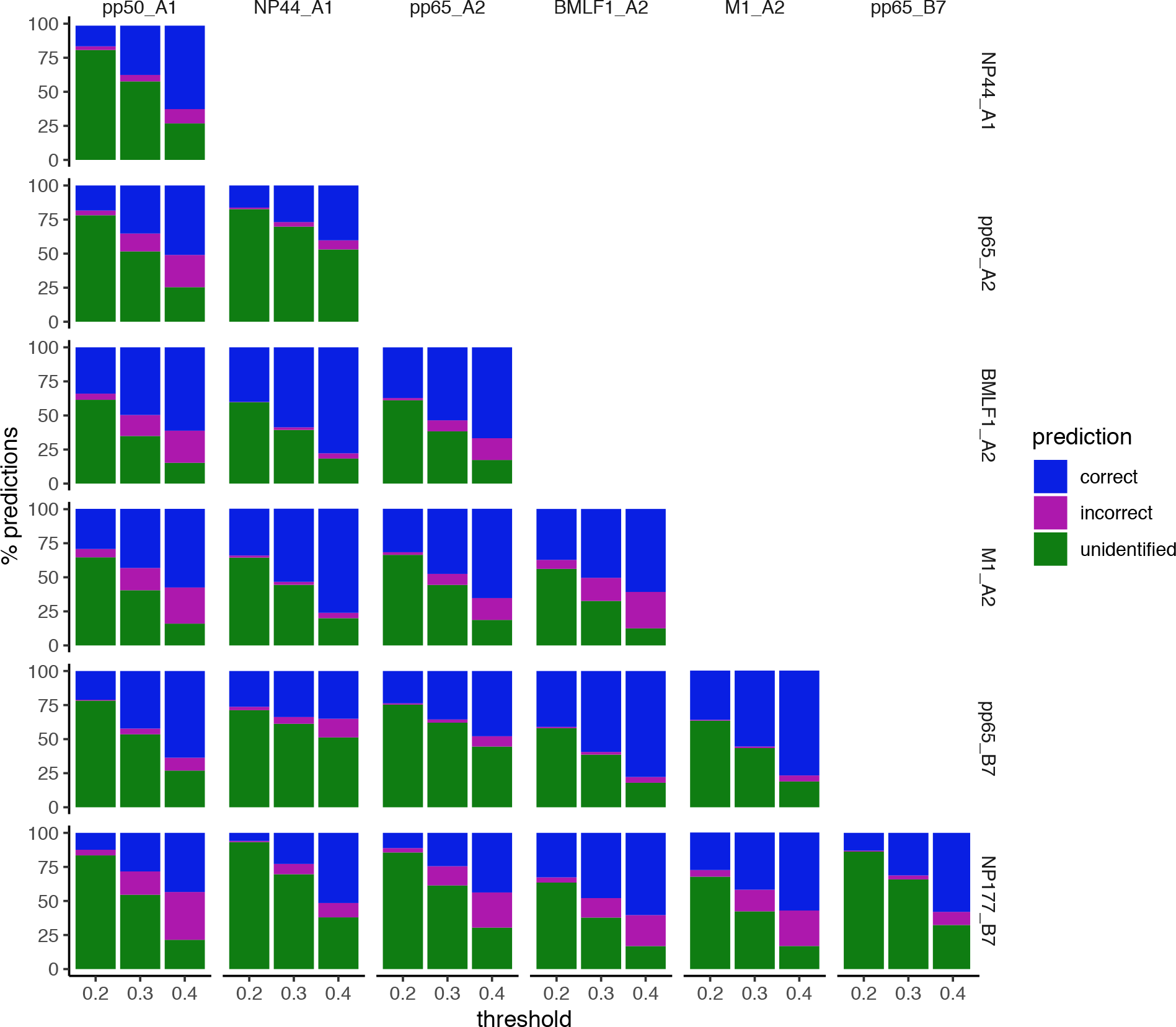
Epitope specificity in Glanville et al. repertoires. Each sub-plot summarizes performance of nearest-neighbor classification, using sequence similarity to predict a CDR’s epitope+HLA from a set of seven different possibilities. The bars indicate the percentage of correct (blue), incorrect (magenta), and unidentified (green) predictions between a pair of epitope+HLA specificities (row and column) at thresholds of 0.2, 0.3, and 0.4.

### Classification of pathology by CDR similarity

McPAS-TCR catalogues TCR sequences from T cells associated with various pathological conditions in humans and mice (53). This repository allowed us to move up from epitope specificity to pathology specificity, evaluating how well CDR3β similarity supports classification of the general pathology from which it was derived. The McPAS-TCR human TCR sets with at least 50 unique CDR sequences were downloaded and split into two groups: “small”, with fewer than 400, and “large” with more than 400, yielding a relatively equal number of repertoires per group with relatively balanced number of sequences per repertoire (Tab. 2). All pairs of pathologies within the same size group were then subjected to nearest-neighbor classification, predicting the pathology of one CDR based on that of the most similar one.

**Table 2.** Pathology-associated repertoires for small (a) and large (b) size-matched groups.

| **Pathology** | **Category** | **Size** |
| --- | --- | --- |
| Allergy | Allergy | 259 |
| Celiac disease (celiac) | Autoimmune | 70 |
| Multiple sclerosis (MS) | Autoimmune | 116 |
| Rheumatoid Arthritis (RA) | Autoimmune | 270 |
| Clear cell renal carcinoma (clearcell) | Cancer | 68 |
| Hepatitis C virus (HepC) | Pathogens | 85 |
| Yellow fever virus (YF) | Pathogens | 179 |

**b)**
| **Pathology** | **Category** | **Size** |
| --- | --- | --- |
| Diabetes Type 1 (diabetes) | Autoimmune | 724 |
| Melanoma | Cancer | 475 |
| Cytomegalovirus (CMV) | Pathogens | 921 |
| Epstein Barr virus (EBV) | Pathogens | 1061 |
| Human immunodeficiency virus (HIV) | Pathogens | 649 |
| Influenza | Pathogens | 2939 |

The pairwise classification performance (numbers of correct, incorrect, and unidentified) was calculated for thresholds of 0.2, 0.3, and 0.4, in the common “sweet spot” of performance across all studies (Fig. 10). On average over all pairwise classifications in the small repertoire group, the percentage of correct classifications increases from 21% at 0.2 to 36% at 0.3 and 52% at 0.4, traded off against 2%, 7%, and 15% incorrect, respectively, as the fraction of identified sequences goes from 26% up to 44% and finally 68%. Likewise, for the pairwise classifications in the large repertoire group, the fraction of correct classifications averages 26% at 0.2, 52% at 0.3, and 66% at 0.4, with corresponding incorrect classification averages of 5%, 14%, and 22% and identified averages of 38%, 61%, and finally 87%. Over all these comparisons, the standard deviations for correct fractions ranges from 6% to 13%, with some pairs clearly much better than others. In general, larger repertoires are able to classify more sequences than small ones, and do so at a higher accuracy, presumably due to simply having a higher probability of containing a sufficiently close sequence. Some infectious diseases such as influenza, yellow fever, HIV, and hepatitis C all do particularly well in the classification task, even against each other. Cancers seem to be confused with autoimmune diseases, and diabetes performs poorly. Further studies are required to ascertain whether differences in classification performance across specific pathologies and general pathology types reflect inherent immunological differences or reveal artifacts in experimental procedures that can perhaps be mitigated with systematic integration methodologies.

**Fig. 10.**
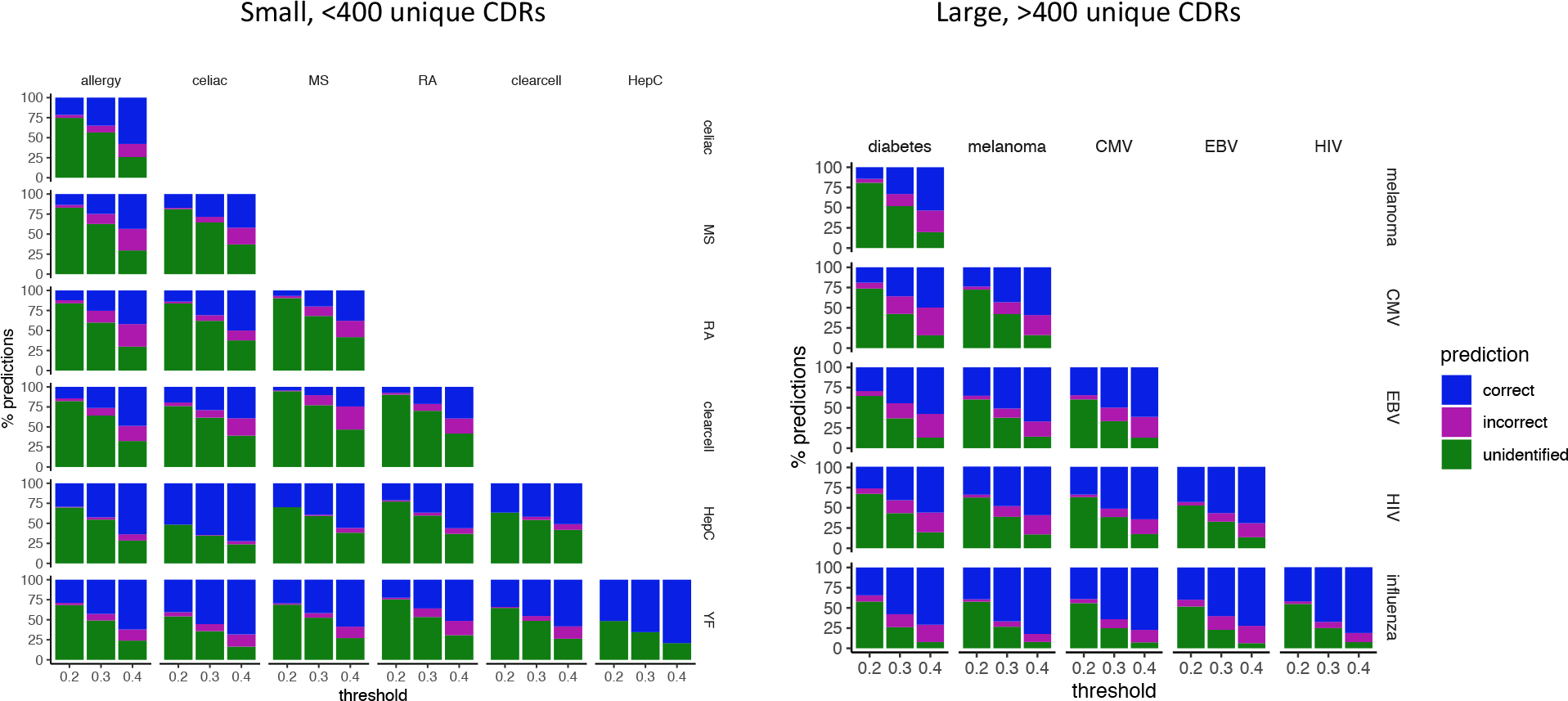
Pathology specificity of McPAS-TCR repertoires. Each sub-plot summarizes performance of nearest-neighbor classification, using sequence similarity to predict a CDR’s pathology association from a set of seven different small repertoires or six different large repertoires. The bars indicate the percentage of correct (blue), incorrect (magenta), and unidentified (green) predictions between a pair of pathologies (row and column) at thresholds of 0.2, 0.3, and 0.4.

## Discussion

This study centers on the use of a representative approach to assessing TCR similarity, with local alignments between size-matched CDR3s for each chain separately, and a relaxed substitution scoring matrix combined with a relatively large gap penalty. This set of choices strikes a balance between looking for the local “hot spots” that mediate binding, and accounting for the overall structural context in which those hot spots are situated. While the detailed outcomes would surely be different if the score moved in one direction toward global alignment or in the other direction toward alignment-free motifs, the same general specificity-sensitivity trends would likely hold. In contrast, integrating information across all six CDRs (and even framework regions) (47–49), rather than considering only CDR3α or CDR3β independently, would likely yield higher overall performance. However, we felt it worthwhile to explore how much information was encoded just in the CDR3 regions, and found them to be strikingly informative.

The 1-nearest-neighbor classifier employed here for specificity-sensitivity assessments is one of the simplest approaches possible, but makes it straightforward to understand and analyze the basis for predictions. A model-based approach, e.g., a linear classifier or even a nonlinear model (50, 51, 54, 55), could give better predictive performance, but would also confound some of the analyses due to the differences in sizes and diversity in different groups. At the same time, a statistical learning approach could provide insights into the importance to different groups of particular CDRs and particular residue positions, directly reveal amino acid motifs conferring specificity, and so forth. In order to focus on the information provided by CDR similarity alone, the analyses presented here did not allow for identity in the 1NN classification and did not consider abundance. This made the classification task somewhat harder, but also provided a uniform basis across the different studies. In a specific practical application, leveraging identity and abundance would be advantageous; statistical learning approaches could offer a natural means to incorporate this information.

Our analysis of structural clusters provided some intriguing insights into sequence-structure relationships, while the rest of the paper explored a range of sequence-function relationships. The structural analysis was somewhat limited due to limited available structures, but a more complete loop modeling approach (56–58) might provide additional power. The functional analysis could benefit from incorporation of MHC restriction information (49), in order to reveal associations among the presented antigens, the presenting MHCs, and the CDRs. And ultimately, combining sequence, structure, and function in an integrated model could provide much deeper insights into the basis for specific recognition.

We individually analyzed repertoires for pairs of twins (46) and for particular antigen specificities (47, 48), along with aggregated collections of pathology-related repertoires (53), but we did not seek to combine information across these different studies. An integrative analysis could provide insights into common modalities of recognition, as has been shown for MHC restrictions (49), but which could also span broader functional associations across antigens from different pathogens as well as from different “self”s. An integrative analysis could thus seek to account for private and public aspects of recognition, gain insights into genetics vs. exposure, and support modeling of the development of immunity.

## Conclusion

This paper has systematically explored the utility of using CDR3 sequence similarity to predict structural class and functional group (namely antigen specificity or pathology association). Based on a representative measure of similarity and an interpretable classification method, the information content in CDR3 alone was shown to support highly specific predictions at a sufficiently stringent similarity threshold, and to maintain good specificity even while increasing sensitivity by substantially relaxing the threshold. Furthermore, patterns supporting predictions within and between groups were shown to provide insights into the structural and functional bases for recognition. We conclude that, if suitably controlled as demonstrated here, predictive frameworks can productively leverage sequence patterns in characterizing and predicting TCR sequence-structure-function relationships.

## Acknowledgments

This work was supported in part by NIH grant 2R01GM098977. We thank Jennifer Franks and Yoonjoo Choi for valuable comments on this work.

## Methods

### Data Processing

When data was processed for each repertoire, the first “C” from each sequence was removed, and duplicate sequences were combined within each repertoire.

### Sequence similarity score

Given a pair of CDRs to evaluate for similarity, local alignment was performed using the Smith-Waterman (SW) algorithm (59), implemented in the Python package swalign (60). SW was applied with the BLOSUM45 substitution matrix (61) to allow for biochemical diversity, and a gap penalty of −10 to focus on matching largely gap-less substrings. Manual inspection of some CDR alignments suggested that these parameters accomplished the intended goals. So as to generate a score that is universally comparable across different CDR sets, the alignment score was normalized by dividing it by the minimum of the self scores of the two sequences. To provide a dissimilarity measure suitable for clustering, the normalized score was then subtracted from 1; since SW scores are non-negative and self-scores are maximal, the final score is between 0 (identical) and 1 (no discernable similarity). Formally, for two sequences *A* and *B*, the distance is:

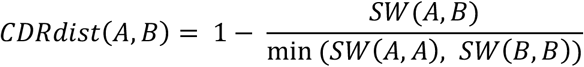

### Nearest neighbor classification

Given a set of CDRs that are labeled as belonging to one of two or more different groups, a nearest neighbor classifier predicts the label of another CDR based on “nearby” labeled CDRs, in terms of the sequence similarity score. A 1-nearest neighbor classifier was used for the results here, making the assignment on the single closest (but not identical) CDR rather than taking a vote among several. Furthermore, the allowed similarity was thresholded, such that if no neighbor was sufficiently similar, then no prediction would be made.

### Clustering

CDRs were clustered using hierarchical agglomerative clustering via the linkage function in the scipy.cluster.hierarchy package of scipy (62), with the sequence similarity score as a comparison function. The resulting dendrogram was cut at a specified sequence similarity threshold in order to define clusters.

### Cluster similarity score

In order to characterize how specific or non-specific clusters were to the repertoires they came from, the distance between a pair of clusters was computed as the average pairwise distance between their members:

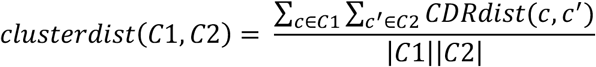

Then the relative specificity of a cluster to its repertoire was characterized in terms of its distance to clusters from other repertoires, with a small score indicating non-specificity (i.e., similarity to a cluster from another repertoire).

### Logos

Sequence logos were generated by Weblogo version 2.8.2 (63, 64) in conjunction with the biopython (65) motif library Bio.motifs.Motif.

